# Micro-indentation and optical coherence tomography for the mechanical characterization of embryos: Experimental setup and measurements on fixed chicken embryos

**DOI:** 10.1101/553693

**Authors:** Marica Marrese, Nelda Antonovaité, Ben K.A. Nelemans, Theodoor H. Smit, Davide Iannuzzi

**Affiliations:** Department of Physics and Astronomy, Laser LaB Amsterdam, Vrije Universiteit Amsterdam, The Netherlands; Department of Orthopaedic Surgery, Amsterdam University Medical Centers, Amsterdam Movement Sciences, Amsterdam, The Netherlands; Developmental Biology, Utrecht University, Utrecht, The Netherlands; Department of Medical Biology, Amsterdam University Medical Centers, Amsterdam, The Netherlands

**Keywords:** mechanical properties, viscoelasticity, micro-indentation, optical coherence tomography, chicken embryos

## Abstract

We introduce an experimental technique that combines micro-indentation and optical coherence tomography to map the viscoelastic properties of embryonic tissue and investigate correlations between local mechanical features and tissue morphology.

**Abstract:** The investigation of the mechanical properties of embryos is expected to provide valuable information on the phenomenology of morphogenesis. It is thus believed that, by mapping the viscoelastic features of an embryo at different stages of growth, it may be possible to shed light on the role of mechanics in embryonic development. To contribute to this field, we present a new instrument that can determine spatiotemporal distributions of mechanical properties of embryos over a wide area and with unprecedented accuracy. The method relies on combining ferrule-top micro-indentation, which provides local measurements of viscoelasticity, with Optical Coherence Tomography, which can reveal changes in tissue morphology and help the user to localize the indentation locations. To prove the working principle, we have collected viscoelasticity maps of fixed HH11-HH12 chicken embryos. Our study highlights the nonlinear behavior of the tissue and qualitatively shows the correlation between local mechanical properties and tissue morphology for different regions of interest.

## Introduction

During embryogenesis, embryos experience a sequence of morphogenetic processes that, under the influence of a complex signaling network, shape the organism and form the organs (Miller and Davidson, 2013a; Wozniak and Chen, 2009). As the mechanical properties of biological tissues are known to influence cell behavior in terms of differentiation, migration, and body formation (Oster et al., 1983; Patwari and Lee, 2008; Varner and Taber, 2012; Varner et al., 2010), one would expect that also the local viscoelastic features of a growing embryo have a strong effect on this process (Mammoto and Ingber, 2010; Miller and Davidson, 2013b; Oster et al., 1983; Patwari and Lee, 2008; Pouille et al., 2009; Varner and Taber, 2012; Wozniak and Chen, 2009). Unfortunately, however, the origin and roles of the forces driving morphogenesis are still in large part unclear. Furthermore, the mechanical properties of embryos are essentially heterogeneous because of the appositional growth of the axial skeleton with the cranial end of the embryo being older and stiffer than the caudal end. It is therefore worth asking whether, from a technical point of view, it is possible to implement novel experimental approaches that could provide new data on the mechanical properties of growing embryos and thus allow the development of more accurate morphogenetic theoretical models.

The mechanical properties of embryonic tissues have already been investigated via a wide variety of techniques, including an uniaxial and compression study (Zhou et al., 2009), microaspiration (von Dassow and Davidson, 2009), macroscopic rheology (Valentine et al., 2005), indentation (Agero et al., 2010; Chevalier et al., 2016; Moore, 1994; von Dassow and Davidson, 2009; Zamir and Taber, 2004a; Zamir and Taber, 2004b) and imaging (Hollnagel et al., 2007; Hove et al., 2003; Jones et al., 2004; Liebling et al., 2006). Uniaxial and compression tests, macroscopic rheology and imaging techniques only measure bulk properties of the embryonic structure and are not suitable to assess the local mechanical response of the different morphological regions of the embryo. Micro-aspiration, which relies on the application of negative pressure on small portions of the embryonic tissue for the evaluation of its elastic modulus, does provide information about the local elastic properties of the tissue, but the technique is highly invasive.

Micro- and nano-indentation setups have in principle the capability to measure local viscoelastic properties without damaging the embryo. So far, these techniques have been mainly used for specific embryonic structures, such as the embryonic chicken heart (Zamir and Taber, 2004a; Zamir and Taber, 2004b) and brain tissue (Xu and Kemp, 2010). Other studies focused on the elastic characterization of cells aggregated from extracted embryonic tissue (Davidson et al., 1999) or on the elastic modulus of embryonic tissue from Xenopus laevis (Adams et al., 1990; Greven et al., 1995; Luu et al., 2011; Moore et al., 1995; Zhou et al., 2009). A few studies have proposed micro- and nano-indentation as a tool to measure the mechanical properties of embryos at specific developmental stages. For instance, in 2010, Agero et al., examined the Young’s modulus of chicken embryos using a micropipette indenter attached to a micromanipulator mounted on an inverted microscope (Agero et al., 2010). These measurements provided new insights concerning the Young’s modulus of the midline (2.4±0.1 kPa), the area pellucida (2.1±0.1 kPa) and the area opaca (11.9±0.8 kPa), but lacked information about the local structures (i.e., somites, presomitic mesoderm, and tail) of the embryonic tissue. More recently, Chevalier and coworkers (Chevalier et al., 2016) used Atomic Force Microscopy (AFM) and a calibrated glass fiber cantilever indenter to assess, respectively, the local and bulk measurements of native (8-day old embryonic midgut) and fixed embryonic tissue, and observed that the elasticity inferred from the AFM measurements was an order of magnitude lower than the one obtained with bulk tests. This discrepancy suggests that bulk tests describe the whole tissue as a composite material, while micro- and nano-indentations are more suitable to extract material properties at local scales. The latter may be more relevant, as the mechanical properties of the micro-environment are known to affect tissue development and change with development. While this study shows that the AFM can provide local and direct mechanical measurements of native (small) embryonic tissue, it seems that also this approach, as all the ones previously discussed, is not capable to investigate possible correlations between mechanics, morphology, and tissue structure at the micro- and mesoscopic scale.

The need to image and mechanically characterize embryonic tissue has driven scientists to use different techniques to image the sample while simultaneously measure its mechanical properties. For instance, in 2015, Filas et al. combined OCT with micro-indentation and numerical analysis to evaluate variations in chicken embryo strain. In this study, a cantilever introduces a small indentation at specific regions, while the OCT system is used to determine the deformation profile near the indentation site from the recorded motions of high-contrast markers injected into the tissue (Filas et al., 2015). However, this method requires an extensive sample preparation and does not provide an absolute measure of the applied force – a piece of information that is necessary for the quantification of the local viscoelastic properties. More recently, two different groups made use of Brillouin spectroscopy to map the mechanical properties of mouse embryos (Raghunathan et al., 2017) and of the zebrafish embryo spinal cord during development and injury (Schlüßler et al., 2018). While this technique allows one to gather mechanical maps that resemble the sample structure, the correlations between Brillouin measurements and stiffness is still ambiguous (Scarcelli and Yun, 2018; Wu et al., 2018). It has been shown, in fact, that water content dominates Brillouin signals, thus providing a measurement of sample compressibility rather than sample stiffness. Moreover, Brillouin scattering is sensitive to gigahertz frequencies, and it does not measure the sample at biologically relevant frequencies (Wu et al., 2018).

From the discussion above, it can be concluded that, despite the numerous studies in the field, there is at present no tool to systematically analyze the local mechanical properties of embryonic tissue, more particular: an adequate instrumental techniques that accurately maps the local viscoelastic response of the embryo at both the micro- and mesoscopic scales.

To solve this impasse, we developed a tool that combines a cantilever-based micro-indentation setup with a non-invasive OCT imaging system to infer the local viscoelastic properties of embryos while simultaneously monitoring its morphological features. To demonstrate the added value of the approach proposed, we performed depth-resolved viscoelasticity maps of chemically fixed HH11-HH12 chicken embryos along the mesoderm from the rostral somite to the caudal tip of the tail. Those maps clearly show the inner morphological and mechanical heterogeneity of three main embryonic regions of interest, from young to old: the tail, the presomitic mesoderm, and the somitic mesoderm. To further validate the potential of the approach proposed, we demonstrate the non-linear behavior of chicken embryos by evaluating its mechanical properties as a function of indentation strain (Antonovaite et al., 2018; Lin et al., 2009a).

## Results

The indenter is based on ferrule-top technology (Chavan et al., 2010; Gruca et al., 2010), where the deflection of a micro-machined cantilever, equipped with a spherical tip, is used to determine the viscoelastic properties of the embryo via depth-controlled oscillatory ramp indentation (Fig. 1A-D) (Antonovaite et al., 2018; van Hoorn et al., 2016). The OCT system images the embryonic structures during the indentation measurements and allows localizing the indentation point and evaluating the quality and the immobilization of the sample. Combining the OCT sections with the indentation curves, one can produce a map that represents the local mechanical properties of the sample.

**Fig. 1.**
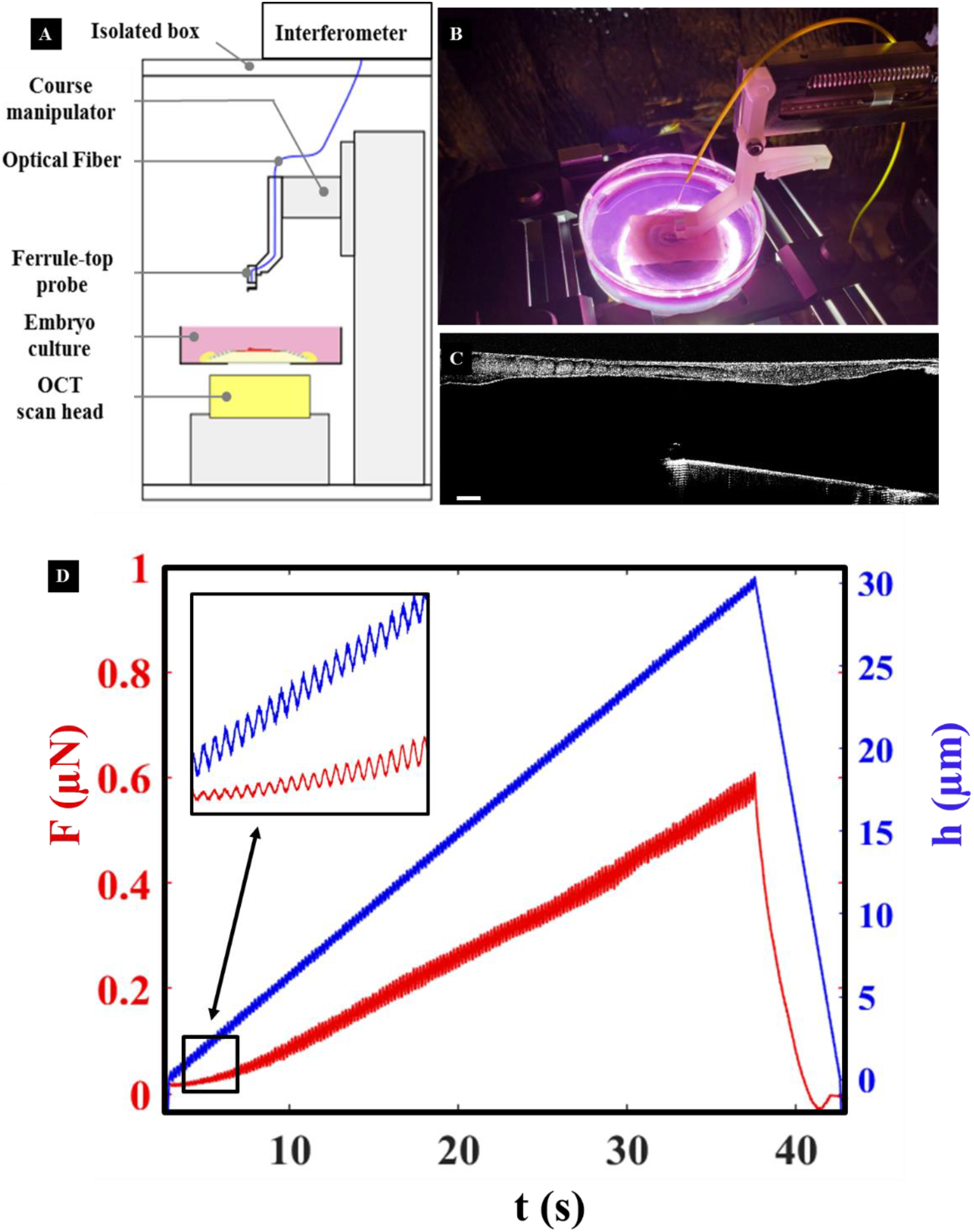
Ferrule-top cantilever setup with OCT for measuring viscoelastic properties of chicken embryos. **(A)** Scheme of the experimental setup. The setup was placed in a wooden box that isolates it from outside vibrations. The box was heated up to 37.7°C before the experiments were started. In the box, the embryo was visualized by OCT from below, while the mechanical measurements were performed by indentations from above. **(B)** Chicken embryo culture with a cantilever probe in the box. **(C)** Sagittal OCT section of a cantilever with sphere, hovering ventrally the chicken embryo. Scale bar 100 μm. Note that the image is upside down because the OCT measures from below the sample. **(D)** A typical depth-controlled indentation oscillatory ramp loading profile. The left y-axis refers to the load; the right y-axis refers to the indentation depth; the x-axis represents time.

For the proof of concept reported in this manuscript, we used HH11-HH12 chicken embryos. All embryos were chemically fixed for 2 or 16 hours and immobilized, after fixation, on an agarose substrate (Fig. 2). The details of the experimental setup and sample preparation are discussed in the Method section.

**Fig. 2.**
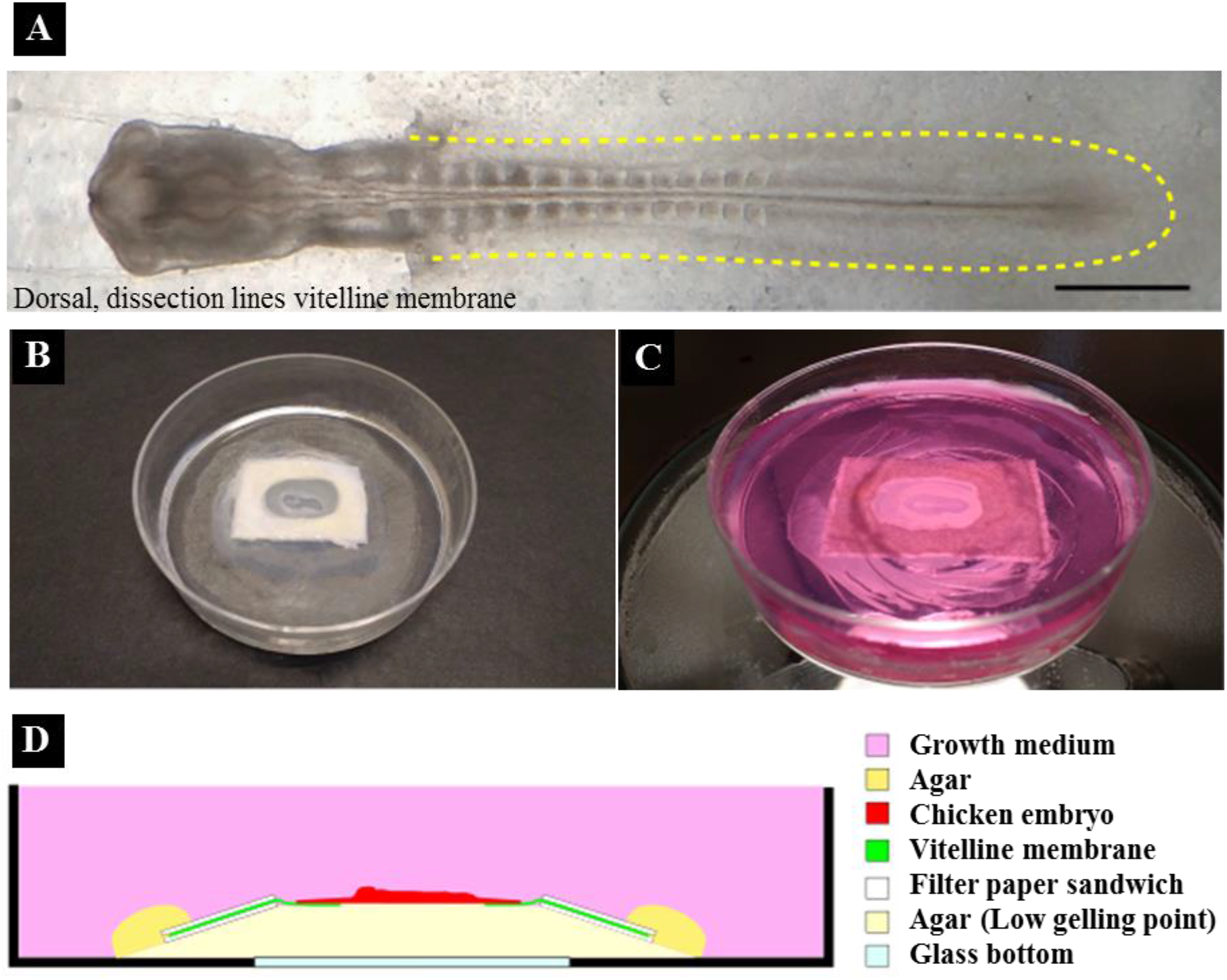
Agarose-immobilized culture for chicken embryos. **(A)** Dorsal view of a HH11 chicken embryo (40hpf). The yellow line indicates the dissection sites where the vitelline membrane is opened along the embryo, to allow the immobilization of the sample in agarose. Scale bar 500 μm. **(B)** The sandwich paper culture chicken embryo is immobilized in agarose by placing the embryo with its dorsal side on low gelling point agarose and covering the filter paper edges with agarose. After the agarose is cured, the embryo is covered with growth medium. **(C)** Side view of embryo culture. **(D)** Scheme showing the agarose immobilization of the chicken embryo. The embryo is embedded in agarose on its dorsal side, while the ventral side is approachable for measurements.

To fully characterize the mechanical properties of the chicken embryos, we performed indentations along the mesoderm from the rostral somites to the caudal tip of the tail (Fig. 3A-D). We also indented transversely so that five regions of interest were measured: lateral mesoderm (LM: regions 1 and 5), paraxial mesoderm (PM: regions 2 and 4) and midline (MD: region 3). For further details regarding the investigated areas, we refer the reader to Table 1. Fig. 4C and D show high resolution (25 μm x 50 μm step size) viscoelasticity maps (*E*′ = storage modulus; *E*″ = loss modulus, at 10% strain and 5.6 Hz frequency, with a sphere radius of 55 μm) of the somitic region of one embryo fixed for 2 hours. The plots seem to confirm that the local mechanical properties of the sample are correlated with somite morphology. From the viscoelasticity maps, one can also distinguish the separation between individual somites – a result that confirms that the indentation method is able to sense structures underneath the endoderm. The scatter plot in Fig. 4E shows the distribution of storage modulus (*E*″) averaged over nine somites (S-XI to S-III) for 5 regions of interest: left and right lateral mesoderm, left and right paraxial mesoderm, and midline. This set of data shows that the somites (left and right) are softer than the midline but stiffer than the lateral mesoderm.

**Fig. 3.**
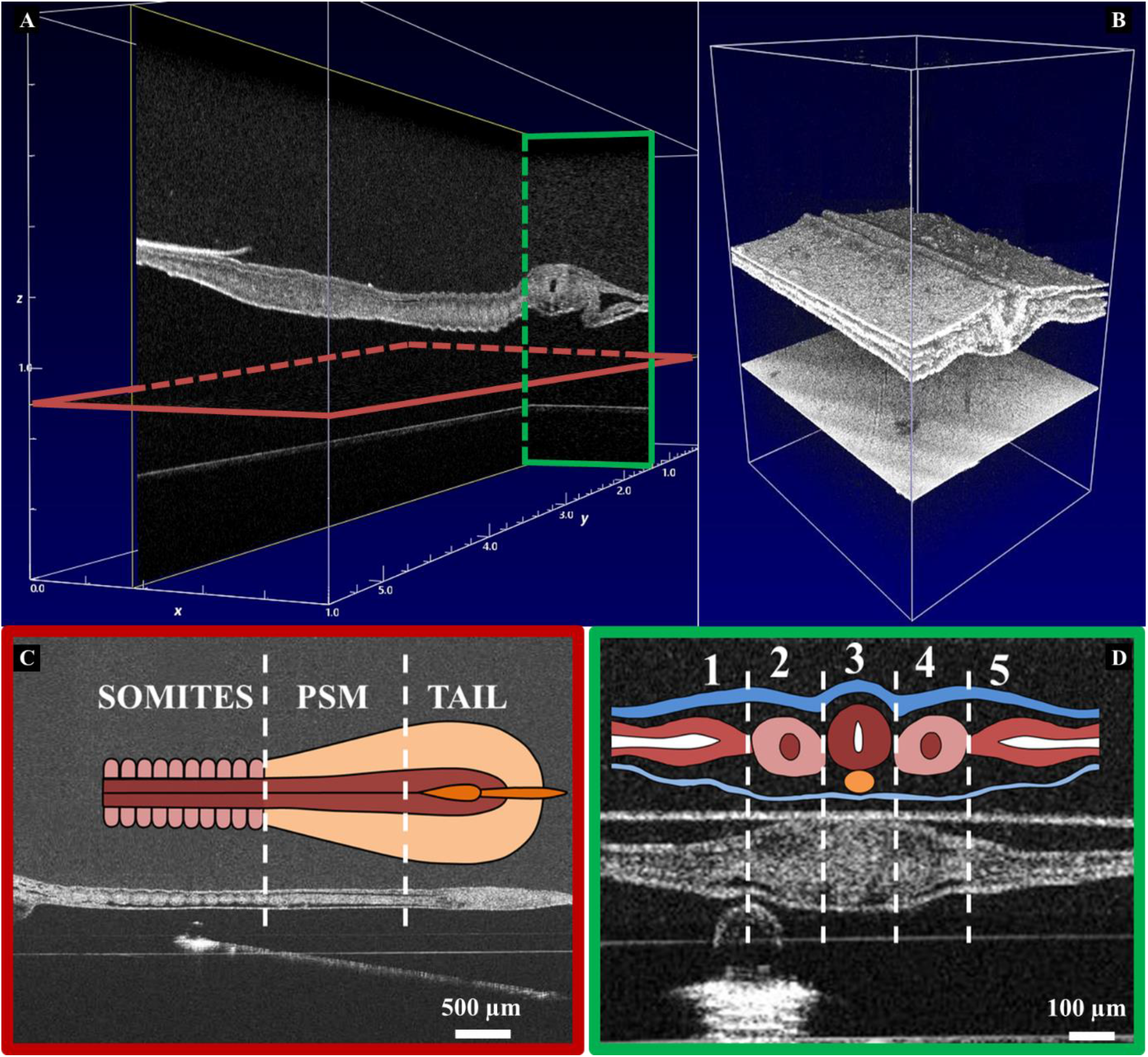
Morphological features of one of the embryo. **(A)** Sagittal (red) and transverse(green) OCT cross section and **(B)** volumetric OCT image showing the three germ layers. **(C-D)** Sagittal and Transverse OCT image of one of the embryo during indentation tests. Note that all the images are upside down because the OCT measures from below the sample.

**Fig. 4.**
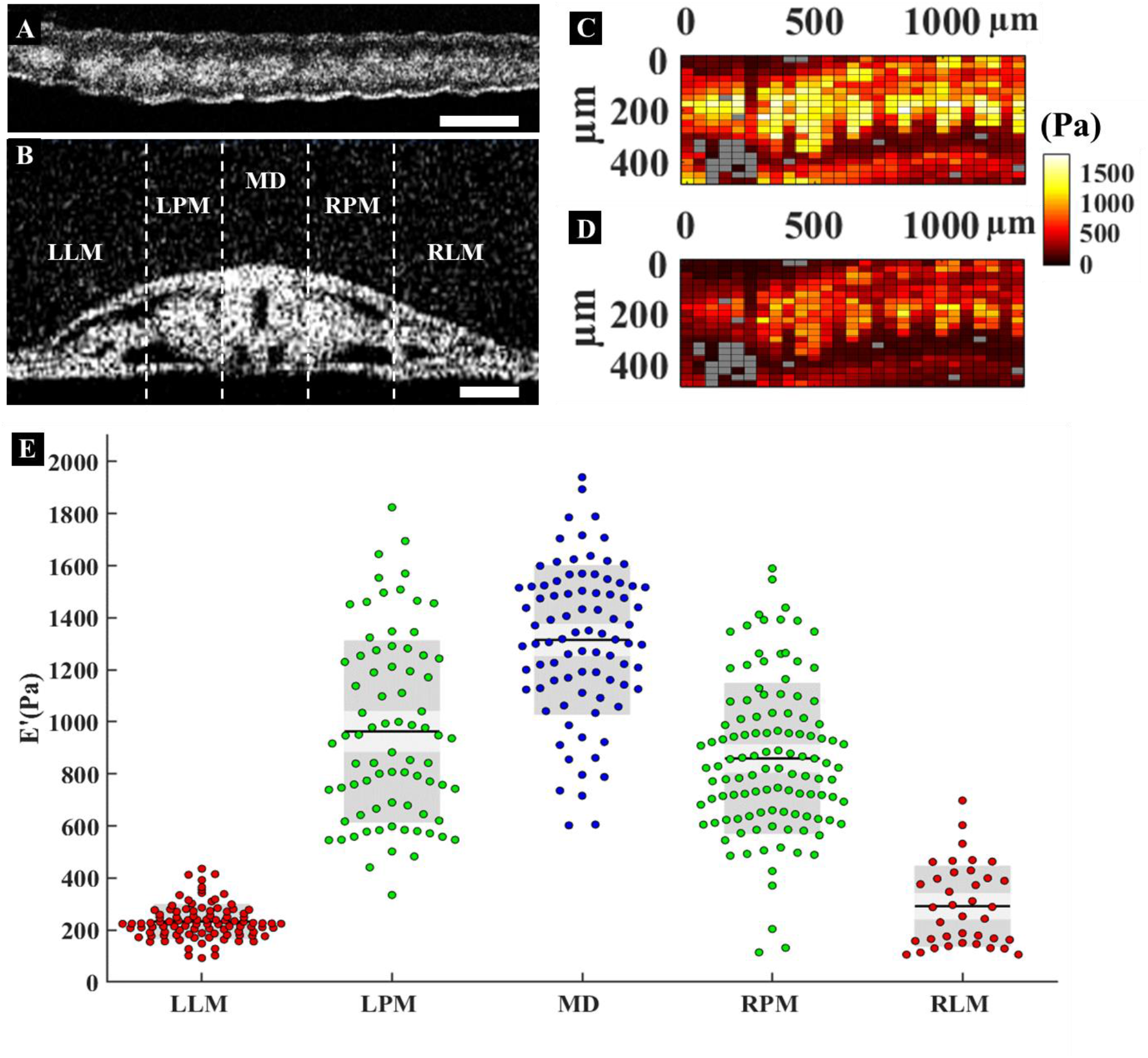
High resolution viscoelasticity map. **(A)** Sagittal OCT section along the somitic region of the embryo. Scale bar = 100 μm **(B)** Transverse OCT image across of the embryo used in this experiment, along with the morphological anatomical regions. Scale bar = 100 μm. **(C-D)** High resolution (25 x 50 μm) colored map of storage *E*′ **(C)** and loss modulus *E*″ **(D)** at 10% strain and 5.6 Hz frequency over the embryo somitic mesoderm. **(E)** Storage (*E*′) modulus across the embryos. Abbreviations of the region of interest from left to right: (LLM) left lateral mesoderm, (LPM) left paraxial mesoderm, (MD) midline, (RPM) right paraxial mesoderm, (RLM) right lateral mesoderm.

**Table 1.**
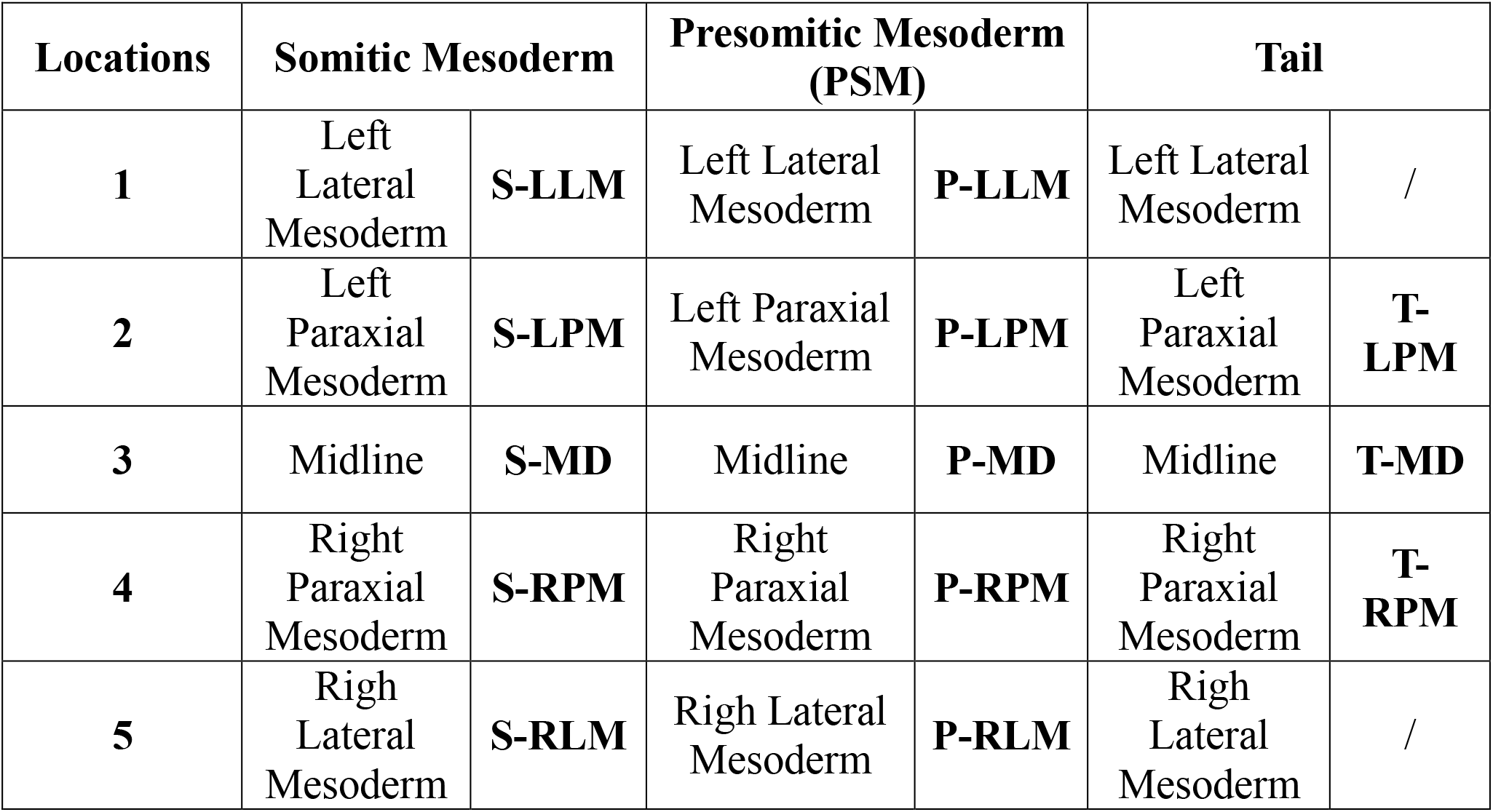
Anatomical regions that were indented.

| Locations | Somitic Mesoderm |  | Presomitic Mesoderm (PSM) |  | Tail |  |
| --- | --- | --- | --- | --- | --- | --- |
| 1 | Left Lateral Mesoderm | S-LLM | Left Lateral Mesoderm | P-LLM | Left Lateral Mesoderm | / |
| 2 | Left Paraxial Mesoderm | S-LPM | Left Paraxial Mesoderm | P-LPM | Left Paraxial Mesoderm | T-LPM |
| 3 | Midline | S-MD | Midline | P-MD | Midline | T-MD |
| 4 | Right Paraxial Mesoderm | S-RPM | Right Paraxial Mesoderm | P-RPM | Right Paraxial Mesoderm | T-RPM |
| 5 | Righ Lateral Mesoderm | S-RLM | Righ Lateral Mesoderm | P-RLM | Righ Lateral Mesoderm | / |

**Table 2.**
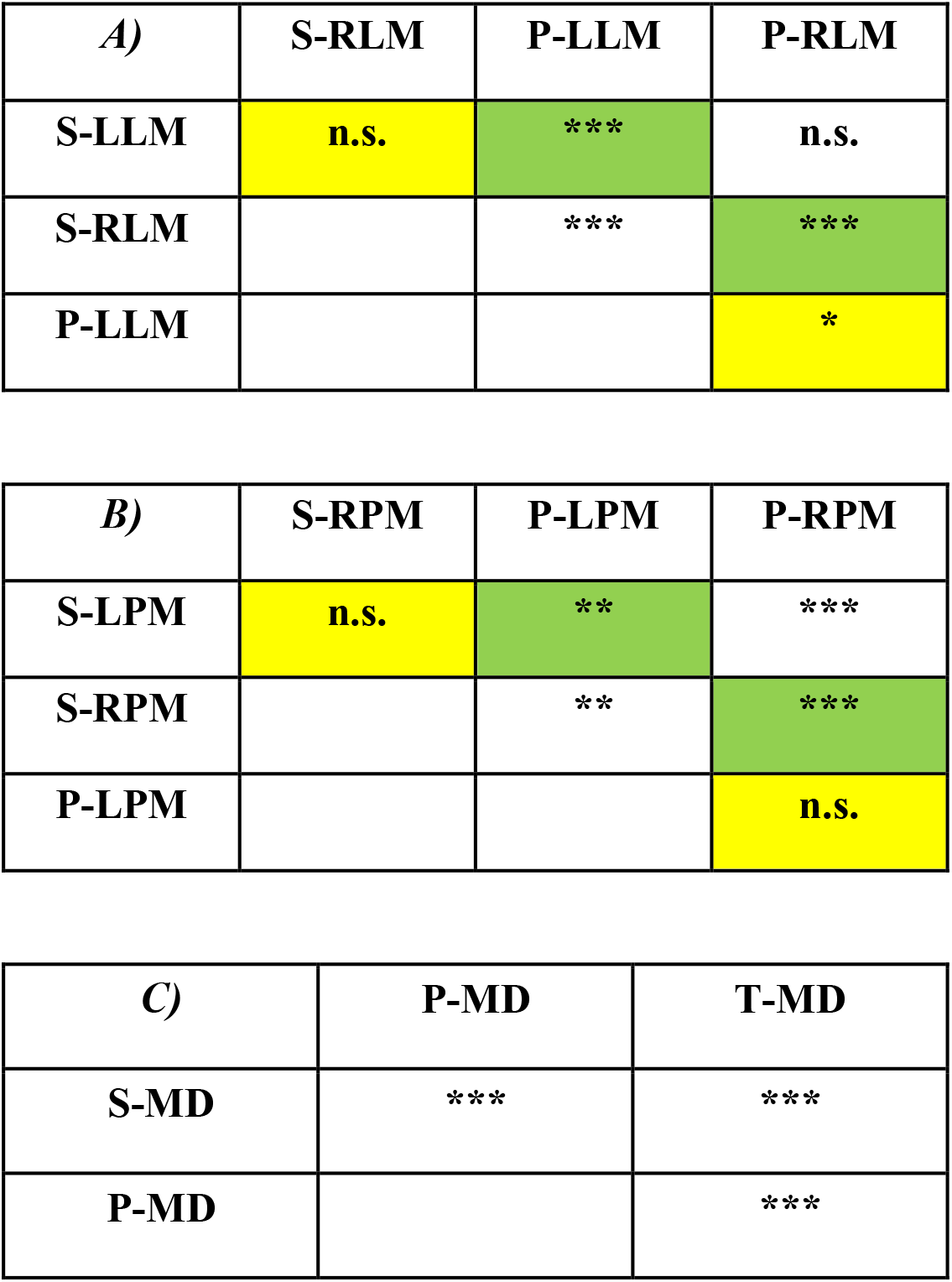
Statistical analysis. Statistical comparison of **A)** lateral mesoderm, **B)** paraxial mesoderm and **C)** midline. Yellow color highlights regions with the same structure on opposite sides of the embryo and green highlight regions with different structures, adjacent to each other.

| **B)** | **S-RPM** | **P-LPM** | **P-RPM** |
| --- | --- | --- | --- |
| **S-LPM** | **n.s.** | **\*\*** | **\*\*\*** |
| **S-RPM** |  | **\*\*** | **\*\*\*** |
| **P-LPM** |  |  | **n.s.** |

| **C)** | **P-MD** | **T-MD** |
| --- | --- | --- |
| **S-MD** | **\*\*\*** | **\*\*\*** |
| **P-MD** |  | **\*\*\*** |

The viscoelasticity map of another embryo (sample fixation time of 16 hours) is shown in Fig. 5A (*E*′) and 5B (*E*″) at 10% strain and frequency of 5.6 Hz covering an area of 800 μm x 4000 μm, from somite VII till the caudal tip of the tail, with a spatial resolution of 50 μm x 50 μm and measured with a sphere radius of 70 μm. Because of the lower resolution and larger indentation sphere used in this second experiment (with respect to the previous one), the mechanical contrast between individual somites is less evident. Yet, this set of data, along with the scatter plot of the storage (*E*′ and loss (*E*″) moduli of the tested sample reported in Fig. 6, still provides valuable information. The mechanical maps together with the scatter plots, in fact, clearly display the morphological heterogeneity of the three main anatomical regions from young to old: tail, presomitic mesoderm and somitic mesoderm. First, the midline is the stiffest structure of the embryonic body, and its stiffness decreases from the somites down to the caudal tip of the tail. The somites are the stiffest material in the paraxial mesoderm, although their difference with the midline does not change significantly while indenting from the somitic region up to the tail. Dynamic indentation reveals that a substantial viscous component is present in the embryonic tissue, as also clear from figures 5B and 6B. Furthermore, elastic and viscous components change along the same trend. Specifically, tan(φ) (see Fig. 7A), which is the ratio between loss and storage modulus, is higher for the paraxial mesoderm and midline and lower for lateral mesoderm in the somitic and presomitic area. This behavior indicates that stiffer and larger structures have higher damping capability and it can also reflect some structural aspect of the tissue. Finally, in the tail, the contribution of elasticity and viscosity is comparable for both the midline and the paraxial mesoderm. While the viscoelasticity results, presented above, were performed at fixed strain and frequency, we also observed that mechanical properties are indentation strain dependent. Fig. 7B, C, and D show storage modulus as a function of the strain for the embryo reported in Fig. 6A. Specifically, all regions stiffen with strain, revealing the non-linear viscoelastic properties of fixed chicken embryos, a result that highlights the importance of measuring the mechanical properties of embryonic tissues in depth-controlled mode (Antonovaite et al., 2018).

**Fig. 5.**
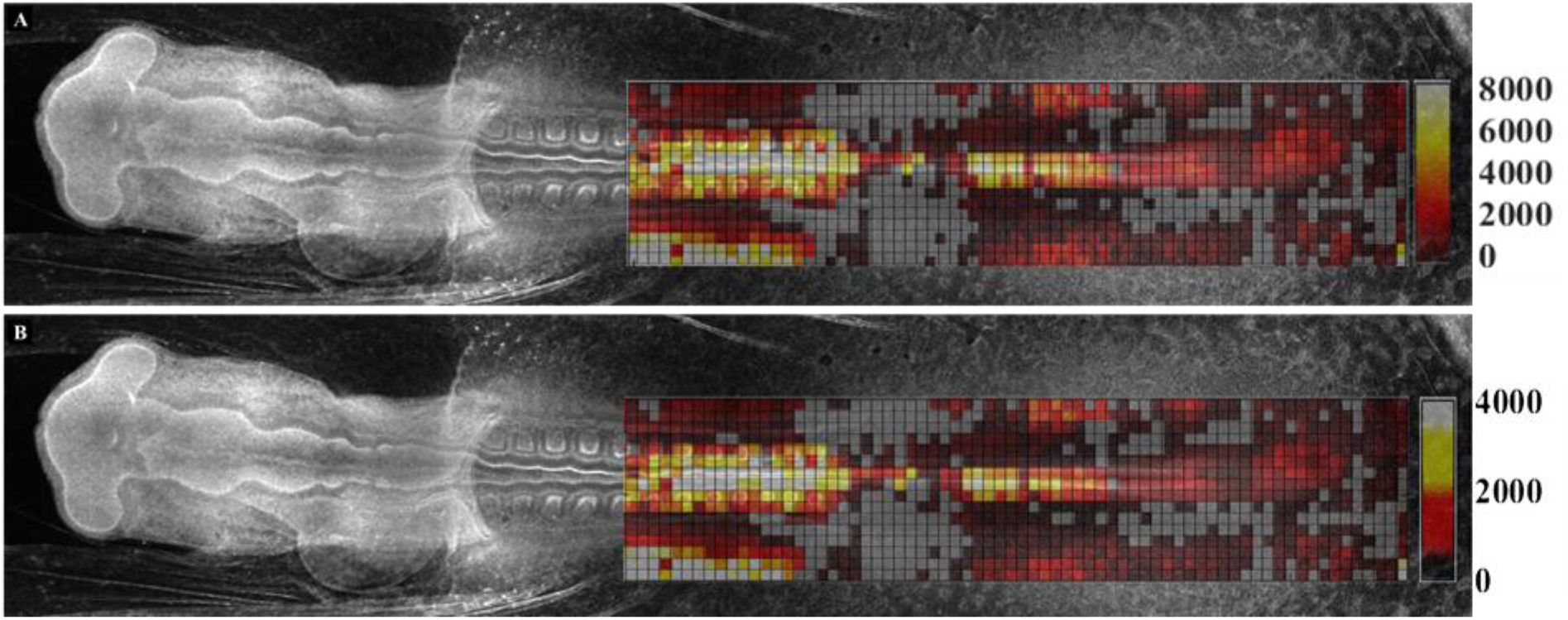
Viscoelasticity maps of one of the chemically fixed embryo. Colored map of storage *E*′ **(A)** and loss modulus *E*″ **(B)** in Pa at 10% strain and 5.6 Hz frequency.

**Fig. 6.**
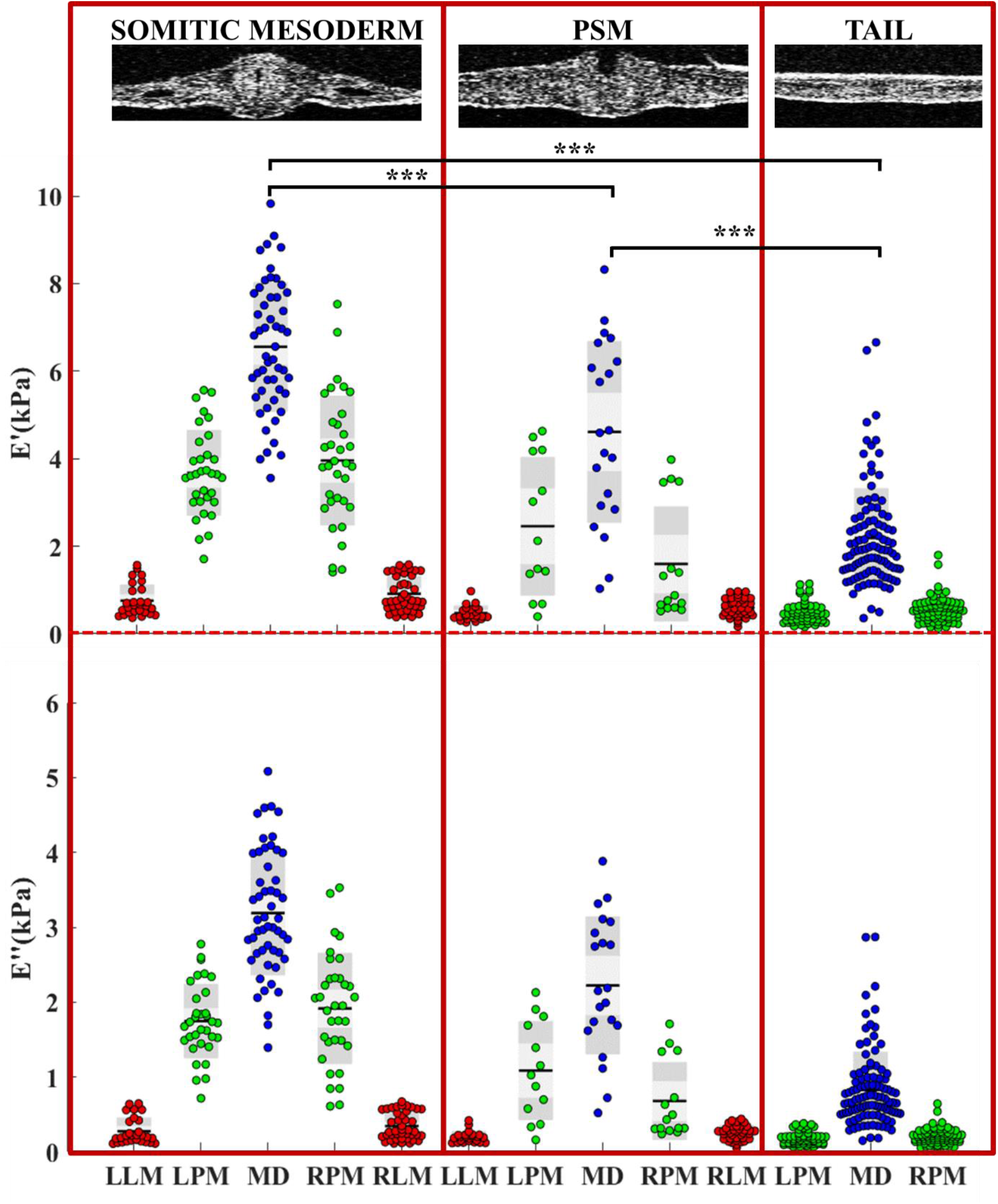
Storage (*E*′) and loss (*E*″) moduli at 10% strain and 5.6 Hz frequency along the embryos. Transverse OCT sections show the three distinct anatomical regions: Somitic Mesoderm, Presomitic Mesoderm (PSM) and Tail. For each region five locations of interest are measured: (LLM) left lateral mesoderm, (LPM) left paraxial mesoderm, (MD) midline, (RPM) right paraxial mesoderm, (RLM) right lateral mesoderm.

**Fig. 7.**
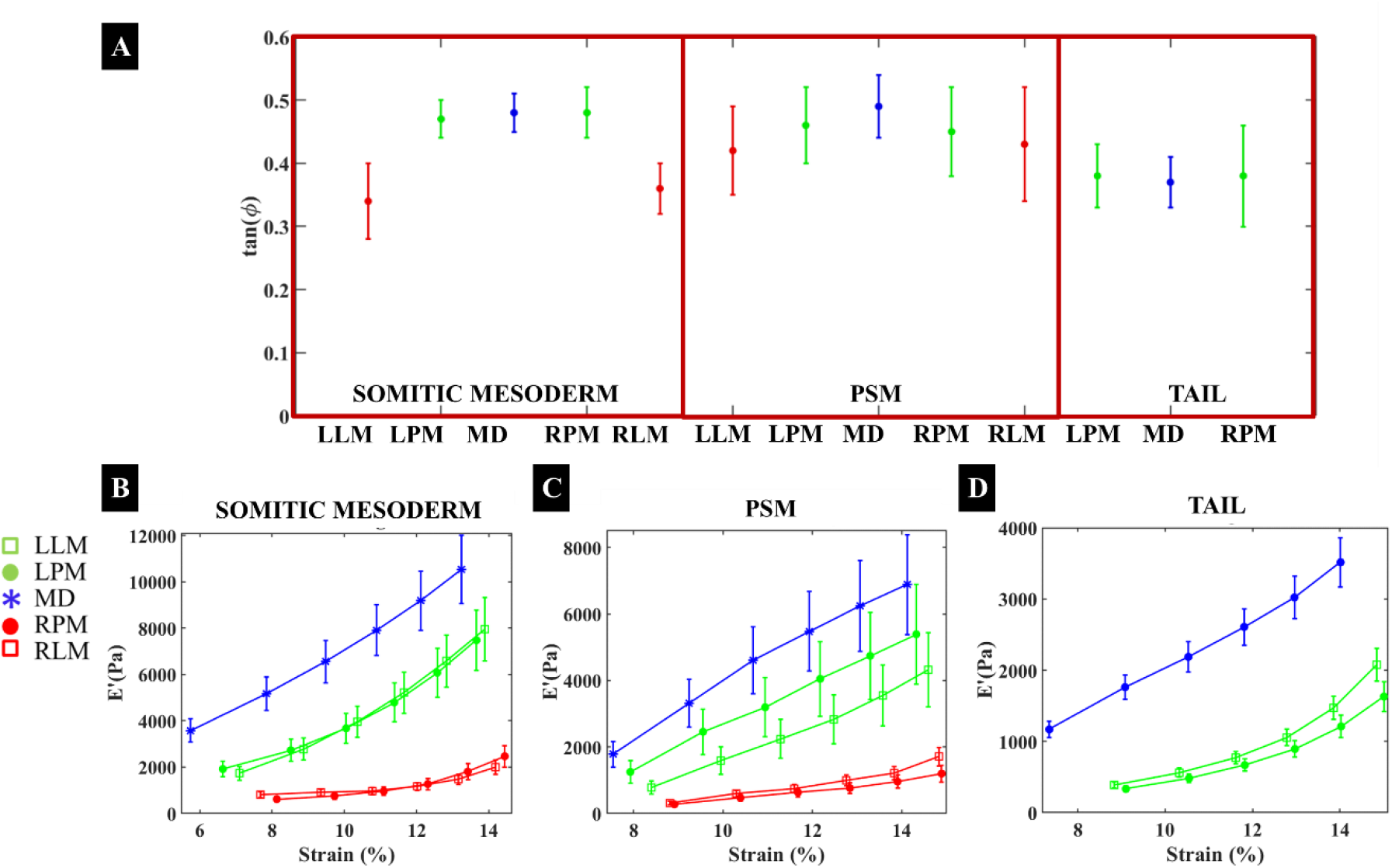
Non-linear and viscoelastic behavior. Loss tangent along the embryo rostrocaudal axis **(A)**. Non-linear viscoelastic properties of chemically fixed embryos obtained with depth-controlled oscillatory ramp indentation. *E*′ increases over the strain range of 6-14% for the somitic mesoderm **(B)**, PSM **(C)** and Tail **(D)**. Note that the standard deviation is positively correlated, since the data are not independent.

## Discussion

To understand the role of mechanical forces in embryogenesis, detailed knowledge of the morphological features of the chicken embryo is crucial. In this context, the use of real-time OCT imaging allows for distinguishing the individual embryonic structures such as somites, notochord, and neural tube. This structural heterogeneity, distinctly visible in the OCT cross-section images (Fig. 3A-D) results in different regional stiffness (Fig.4). It is interesting to note that, even if the sample was chemically fixed, some of the mechanical features observed correlate well with what one would expect for a not fixed sample. For example, the midline, the stiffest structure of the embryonic tissues, contains an embryonic cellular rod, the notochord, that gives structural support to the adjacent tissues, thus representing the major skeleton element in the embryonic vertebrae (Corallo et al., 2015). The somites, the precursors of the vertebrae, cartilage, tendons, and dermis, is softer than the midline but stiffer than the lateral mesoderm, which mostly forms the circulatory system, body cavity, and pelvis in the adult body (Gilbert, 2013). Thus, we may assert that some of the spatial variations in stiffness are related to the physiological roles of each anatomical structure.

From the analysis of the indentation maps in Fig. 5 along with OCT images, one can observe a consistent correspondence between the morphology of the three indented regions and their mechanical properties. These indicate that a gradient of stiffness can be identified along the mesoderm from the rostral somites to the caudal tip of the tail. Once again, it seems that the fixed embryo resembles mechanical behaviors that would fit well within a description of the mechanical properties of a live sample. This trend, in fact, could be correlated with the maturation of the PSM. The somites are formed in pairs of epithelial spheres beside the neural tube, whereas in the PSM the somites are not entirely epithelialized yet. The stiffness along the body axis could also reflect the maturation of each structure, in fact, at this stage, the cranial end of the embryo is older and stiffer than the caudal end. If this behavior is confirmed *in vivo*, one could argue that the biomechanical processes during somite formation, such as changes in extracellular matrix composition, migration, and contraction of mesodermal cells, could be triggered by this gradient of stiffness along the rostrocaudal axis of the embryo. However, structural components responsible for different viscoelastic properties should be investigated in more detail by studying the embryonic tissue *in vivo*.

In conclusion, we have introduced a new tool that is able to accurately capture variations in stiffness due to different sample structure or treatment, to reliably measure the viscoelastic and non-linear properties of heterogeneous materials and, simultaneously investigate sample morphology.

Specifically, our findings, suggest that for each experiment, the variation in terms of absolute stiffness values could be strictly related to the sample treatment (chemical fixation time). The embryo that was used to produce the data of Fig. 4, in fact, was fixed for only 2 hours while, for Fig. 5, we used a sample with much longer fixation time (16 hours), which give rise to a stiffer material (Kim et al., 2017; Ling et al., 2016; Thavarajah et al., 2012).

By performing depth-controlled frequency-domain indentation tests on embryonic tissues, we have evaluated the viscoelastic nature of the embryonic tissue, as previously observed in time domain with stress relaxation, creep test, and with non-linear finite element modeling for the heart tube in other experiments (Forgacs et al., 1998; Yao et al., 2012). Our viscoelasticity maps further reveal that the viscous component is not negligible for all the indented regions, thus indicating that it is necessary to mechanically characterize the embryonic tissue by considering both the elastic and viscous contribution.

In addition, we have demonstrated that our technique can also reveal the presence of non-linear mechanical behavior of tissues. Both storage and loss modulus increase with indentation strain, thus proving that fixed chicken embryos are a non-linear material, that, to our knowledge, it was never reported in the literature before. This last finding highlights that the mechanical properties of fixed chicken embryos strongly depend on the indentation strain at which each measurement is performed. It is reasonable to assume that a similar behavior can be observed in live embryo, suggesting that, to obtain reliable measurements of the mechanical properties of embryos, it is crucial to rely on a depth-controlled indentation mode.

In the future, the integration of OCT imaging with indentation techniques could be further exploited by introducing mathematical models that provide a more extensive evaluation of different deformation profiles in relation to the local material composition. One of the main limitations of our indentation setup is, in fact, the mathematical model used to extrapolate mechanical properties, as we assume that the sample is isotropic, homogeneous, flat, nonadhesive and that the time-dependent properties are due only to the viscoelasticity, by neglecting the poro-elasticity. Clearly, the developing embryo is none of this. In this perspective, the combination of OCT imaging with indentation tests could potentially be used to model the mechanical properties of heterogeneous tissue by monitoring the deformation of individual structures, as in Optical Coherence Elastography (Kennedy et al., 2015). Moreover, the OCT image could be also used to thoroughly investigate the evolution of tissue mechanical properties during morphogenesis and development and to determine quantitatively mechanical changes in normal and pathological conditions, filling the existing gaps in the field of developmental biology.

## Materials and Methods

### Experimental setup

The setup consists of a cantilever-based indentation arm, an OCT imaging system, and a sample holder (Fig. 1A). The indentation arm comprises of an XYZ micromanipulator (PatchStar, Scientifica, UK), a piezoelectric transducer (PI p-603.5S2, Physik Instrumente) and a ferrule-top cantilever indentation probe connected to an interferometer (OP1550, Optics11, The Netherlands). A single mode optical fiber is used to readout the bending of the cantilever, as previously described (Van Hoorn et al., 2016). Cantilevers with 0.25 and 0.75 N/m spring constant and spheres radius of 55 and 70 μm were used (Fig. 1C), for the experiments performed on the 2 hours and the 16 hours fixed samples, respectively. The radius was chosen to be large enough for indenting deep into the tissue without any slipping effects and small enough to keep a high spatial resolution between somites. The system is placed on a vibration isolation table (1VIS22-60-4, Standa, Lithuania) and enclosed in a custom-made wooden box with acoustic foam covering inside to minimize external noise.

### Optical coherence tomography (OCT)

To find anatomical locations and follow each indentation experiment, the embryos were scanned with a spectral domain SD-OCT (Telesto II series, Thorlabs GmbH, Germany). A super-luminescent diode (SLD, D-1300 HP, Superlum, Ireland) with full width half maxima (FWHM) of 85 nm and central wavelength of 1310 nm was used as light source. This OCT system can provide real-time imaging with maximum imaging depth of 3.5 mm, axial resolution of 5.5 μm in air and 4.2 μm in water, and a transverse resolution of 11.8 μm. The setup was operated in inverted mode as the images are taken from beneath the glass surface of the Petri dish of the agarose culture. Afterward, the collected OCT data were processed using commercial software (ThorImage OCT 4.3, Thorlabs GmbH). All the scale bars are expressed in liquid optical path length (*n*=1.33), unless otherwise mentioned. For all experiments, a B-scan (transverse) image along the left-right embryo axis was acquired to evaluate the quality and the attachment of the sample. Furthermore, during indentation measurements, OCT cross sections were captured every 10 seconds to precisely discriminate each location in the regions of interest (Fig. 3 C and D). Combining the OCT sections and the indentation curves, each indentation was evaluated, and failed measurements were removed according to a predefined scheme.

### Indentation protocol and data analysis

Indentations were performed in a depth-controlled mode by using a feedback loop (van Hoorn et al., 2016). The oscillatory ramp profile (Fig. 1 D) was selected for the characterization of viscoelastic properties at different depths, as shown by Antonovaite *et al*. (Antonovaite et al., 2018). Strain rate of the ramp was 0.01, maximum depth 25 μm, and amplitude and frequency of oscillations were 0.25 μm and 5.62 Hz, respectively. Load-indentation data were used to extract storage and loss moduli by fitting every 20 oscillations to a cosine function (custom written code in Matlab). Fits with squared correlation coefficient *R*^2^ < 0.5 were removed from the data set. Storage modulus values at ~10% strain (corresponding to a depth of ~16.3 μm) were selected for regional comparisons, accomplishing the requirements of *h* < 10% of the sample thickness but not fulfilling the small strain approximation (Lin et al., 2009). From these raw data, the storage and loss moduli (*E*′ and *E*″) were deduced by:

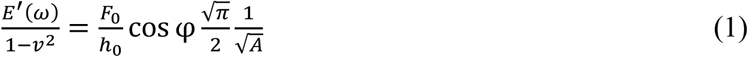

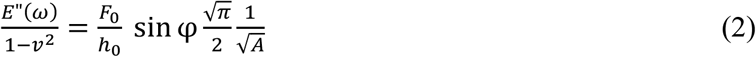

where *E*′ is the storage modulus, *E*″ is the loss modulus, *ω* is the frequency, *F_0_* is the amplitude of the oscillatory-load, *h_0_* is the amplitude of the oscillatory-indentation, *v* is Poisson’s ratio of compressibility (0.5, assuming incompressibility), *φ* is the phase shift between the indentation and load oscillations, *A* = π*a*^2^ is the contact area 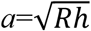, where *R* is radius of the sphere and *h* – indentation depth (Lin et al., 2009b). With the oscillatory ramp, we can define the ratio between the loss and storage modulus, known as loss tangent, *tan(φ)*, which provides the relation between the viscous and elastic components.

### Statistics

Normality of data distribution was tested with Shapiro-Wilk test. In case of normal distribution, statistical differences between test samples were investigated with two-way ANOVA. For non-normally distributed data, non-parametric Kruskal-Wallis ANOVA test was used to compare data samples. All statistical analyses were performed with Statistics and Machine Learning Toolbox (version 2017a, The Mathworks, Natick, MA, USA).

### Chicken embryo cultures

Fertilized chicken eggs, white leghorns, *Gallus gallus domesticus* (Linnaeus, 1758), were obtained from Drost B.V. (Loosdrecht, The Netherlands), incubated at 37.5 °C in a moist atmosphere, and automatically turned every hour. After incubation for approximately 41h, HH11-HH12 chicken embryos (Hamburger and Hamilton, 1992) were explanted using filter paper carriers (Chapman et al., 2001) cultured *ex ovo* as modified submerged filter paper sandwiches (Schmitz et al., 2016), chemically fixed, immobilized in agarose and immersed in growth medium (Palmeirim et al., 1997).

Chicken embryos were explanted as submerged filter paper sandwiches (Schmitz et al., 2016) and washed in Dulbecco’s PBS (Sigma, ref. D8357). On the dorsal side of the embryo, we made a rostrocaudal slit in the vitelline membrane from the heart to the tip of the tail (Fig. 2A) to assure a properly immobilize of the embryo on agarose. After, the chicken embryos were first chemically fixed using 4% of formaldehyde buffered solution (Sigma, ref. 1004965000) for 2 or 16 hours at 4 °C and then washed twice with PBS, before immobilization on agarose. To prepare the embryo culture, 60 mm × 15 mm Petri dishes (Sigma, ref. P5481) were equipped with a glass bottom (30 mm circular cover glass #1, Thermo Scientific Menzel ref. CBAD00300RA140MNZ#0) to allow the OCT to image from below. A solution of 1.5% w/v low gelling temperature agarose (LGT agarose, Sigma, ref. A9414), was kept at 40°C on the bench. A hill of 500 μl 1.5% w/v agarose (Sigma, ref. A9539) was made on top of the glass bottom (Fig. 2F). Then, fresh LGT agarose was pipetted over the agarose hill. The embryo filter paper sandwich was dried horizontally with tissue and then placed on the fresh LGT agarose, dorsal side down. Carefully, the filter paper sandwich was moved horizontally, to allow the penetration of the LGT agarose through the previously made slit in the vitelline membrane and to cure towards the ectoderm of the embryo (Fig. 2B). Because of the hill of agarose, the embryo was pointed upwards, which helped to approach the embryo with the indenter. The edges of the filter paper were covered with agarose to completely immobilize the filter paper. To prevent dehydration of the embryo during the LGT agarose curing, a droplet of medium was carefully brought on top of the embryo, without touching the curing agarose. After approximately 3 minutes, the culture was placed in the indentation box (Fig. 1B) and growth medium was slowly poured into the Petri dish (Fig. 2C). After, the embryo was submerged in 25 ml of the growth medium and anatomically aligned under the OCT scan head. The growth medium consisted of medium 199 GlutaMax (Invitrogen, ref. 41150-020; 4°C), 10% chicken serum (GIBCO, ref. 16110-082; −20°C), 5% dialyzed fetal bovine serum (FBS) (GIBCO ref. 26400-036; −20°C) and 1% of a 10000 U/ml stock solution of Penicillin/Streptomycin (GIBCO ref. 15140-122; −20°C).

## Acknowledgments

The authors acknowledge Erik Paardekam for his technical support.

## Competing Interests

DI declares potential conflict of interest as founder, shareholder, and advisor of Optics11.

## Funding

This work has been financially supported by the European Research Council under the European Union’s Seventh Framework Programme (FP/20072013)/ERC grant agreement no. [615170]. BN was financially supported by ZonMW-VICI grant 918.11.635 to TS.

## Data availability

All raw and processed data of this study are available on reasonable request from the corresponding author.

